# Draft mitochondrial genome of the tree species *Neltuma alba* (Caesalpinioideae, tribe *Mimoseae*) from the Atacama Desert, Chile

**DOI:** 10.64898/2026.01.06.697924

**Authors:** Irina Rojas-Jopia, Mariel Mamani-Gómez, Valeska Rozas-Lazcano, Matias Talamilla-Marivil, Roberto Contreras-Díaz

## Abstract

In this study, we assembled the draft mitochondrial genome of *Neltuma alba*, a vulnerable native species inhabiting the Atacama Desert from Chile. Here, we describe the structure and phylogeny of the complete mitochondrial genome of *N. alba*. The mitochondrial genome consisted of three linear DNA sequences. The whole mitochondrial genome (mitogenome) sequences comprised 717,588 bp in length and contained 36 protein-coding genes, 19 tRNA and three rRNA. A maximum-likelihood phylogenetic analysis based on the seven mitochondrial genomes currently available in NCBI GenBank confirmed the placement of *N. alba* within the *Neltuma* genus, supporting its classification within the tribe *Mimoseae*. This study presents, for the first time, the complete mitochondrial genome of *N. alba*, offering a valuable resource for tracing the evolutionary and geographic diversification of the *Neltuma* genus in America.

## Introduction

Legumes are globally distributed and ecologically essential, particularly in arid ecosystems such as the Atacama Desert. The Fabaceae family, one of the largest among angiosperms, previously included the genus *Prosopis* L., which encompassed 57 species across arid regions of the Americas, Africa, and Asia (Burkart, 1976). Within this group, species from the *Algarobia* section, such as *Prosopis alba* (Griseb.) C.E. Hughes & G.P. Lewis, are drought-tolerant trees capable of thriving in extreme environments, including the hyper-arid Atacama Desert, one of the driest regions on Earth (Garrido et al., 2020). Native to South America, *P. alba* is highly valued for its ecological resilience and multifunctional uses. In the Atacama, it endures intense solar radiation and severe water scarcity, while providing essential ecosystem services to local communities, including food, forage, and timber (Contreras et al., 2020). Recent molecular phylogenetic analyses based on chloroplast and nuclear DNA have demonstrated that *Prosopis* is polyphyletic, leading to a taxonomic revision. Consequently, species from the *Algarobia* section have been reassigned to the genus *Neltuma*, and *P. alba* is now recognized as *Neltuma alba* (Griseb.) C.E. Hughes & G.P. Lewis (Hughes et al., 2022). This reclassification better reflects the evolutionary history of these desert-adapted legumes and highlights their distinct lineage within *Fabaceae*, where the former “*Mimosoid* clade” is now recognized as the tribe *Mimoseae* (Bruneau et al., 2024).

Plant mitochondrial genomes offer valuable insights into evolutionary relationships, aiding taxonomic revisions and revealing lineage-specific adaptations to arid environments. Mitochondria play a central role in energy metabolism and stress response, thereby influencing plant survival under extreme conditions (van Aken, 2021). Despite their complex structure, the conserved coding regions of plant mitochondrial genomes serve as informative molecular markers for phylogenetic inference (Contreras-Díaz et al., 2023). Although often depicted as simple circles, plant mitochondrial genomes exhibit complex structures comprising a dynamic mixture of linear, branched, and circular molecules— reflecting multiple structural isoform (Kozik et al., 2019; Štorchová & Krüger, 2024), including multipartite mitochondrial configurations (Sloan, 2013). Evolutionarily, these genomes are characterized by low mutation rates but high structural variability and frequent recombination, shedding light on genome evolution and hybridization events (Gualberto & Newton, 2017). In this context, we employed next-generation sequencing (NGS) to assemble the mitochondrial genome of *N. alba*, with the aim of elucidating its genome structure and phylogenetic relationships in comparison with other members of the tribe *Mimoseae*.

## Materials and Methods

Fresh leaves of *N. alba* were collected by Roberto Contreras from near Copiapó of Atacama region in Chile (27°21’39.3”S 70°20’33.8”W) (Figure 1). The taxonomic identification of the species was conducted based on the descriptions provided by Burkart et al (1976). Moreover, the voucher specimens (EIF13329) were deposited in the Herbarium EIF of the Facultad de Ciencias Forestales y de la Conservación de la Naturaleza (https://sweetgum.nybg.org/science/ih/herbarium-details/?irn=126262), Universidad de Chile (Nicolás García).

**Figure 1.**
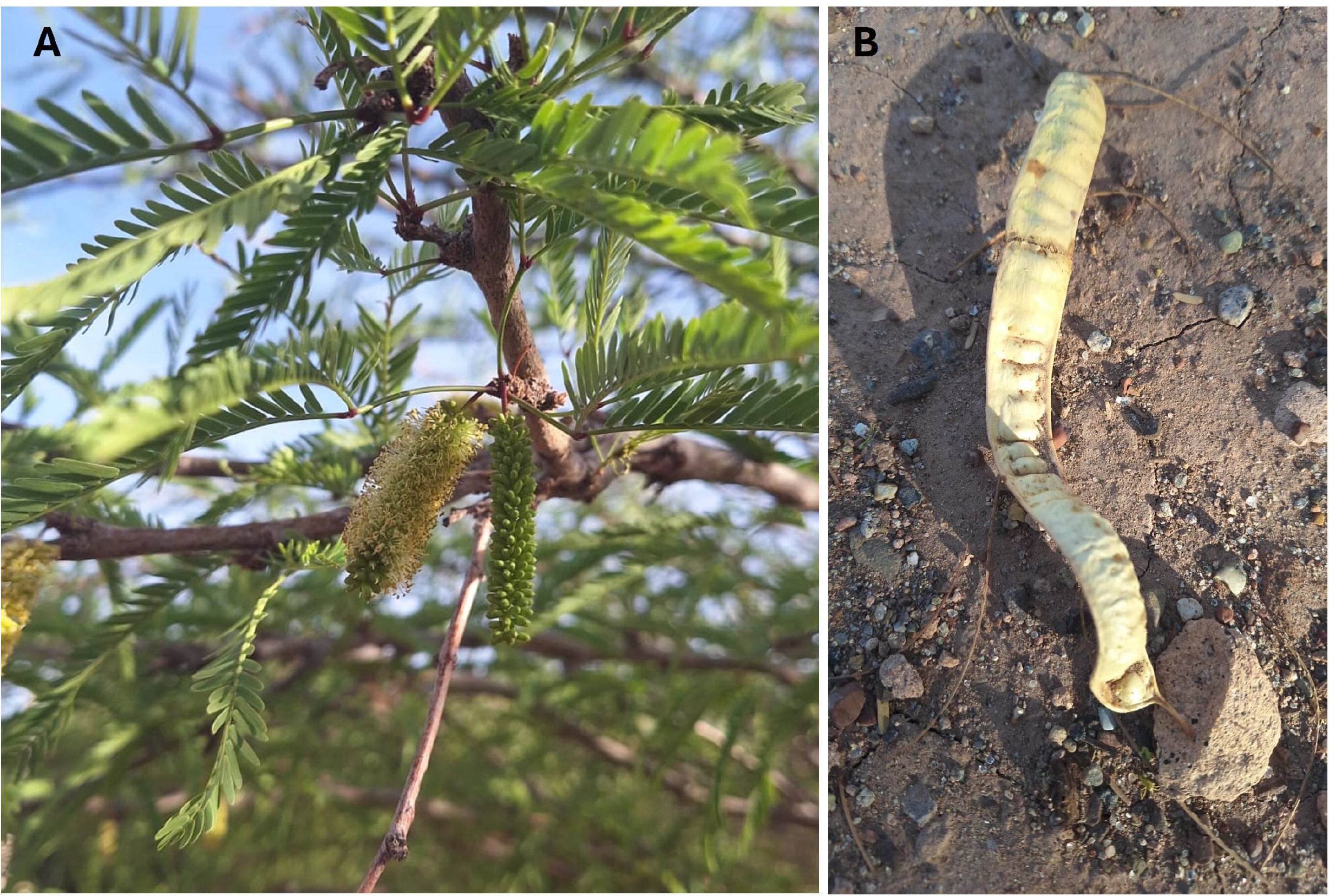
*Neltuma alba*. (A) Leaves and flowers. (B) Fruit. Photographs taken by Roberto Contreras-Díaz.

The total DNA of *N. alba* was extracted using a modified CTAB method (Contreras-Díaz et al., 2025). Paired-end genomic sequencing (PE150) was carried out using the Illumina NovaSeq 6000 platform, yielding approximately 36□GB of high-quality clean reads. Trim Galore (Krueger, 2019) was used to filter low-quality reads and sequencing adapters. To assemble the draft *N. alba* mitochondrial genome (mitogenome), we followed a four-step approach. First, the reads were quality-checked and assembled using MEGAHIT v1.0 (D. Li et al., 2015). Second, we employed GetOrganelle (Jin et al., 2020) to generate additional contigs, using accessions of *Neltuma glandulosa* (Torr.) C.E. Hughes & G.P. Lewis (MW448450, MW448451, MW448452, MW448453, MW448454, and MW448455) as references (Choi et al., 2021). The following command was used as an example: get_organelle_from_reads.py -s MW448450.fasta --genes MW448450-genes.fasta -1 forward.fq -2 reverse.fq -R 8 -k 21,45,65,85,105 -F anonym -o N.alba_mt_out. Third, the contigs generated by both MEGAHIT and GetOrganelle were filtered using SeqKit (Shen et al., 2024) to retain only mitochondrial sequences longer than 1,000 bp, and chloroplast sequences exceeding 1,000 bp were removed. Finally, all filtered contigs were imported into Geneious Prime 2023.2 (https://www.geneious.com/) (Kearse et al., 2012) to assemble the final *N. alba* mitogenome. The complete mitogenome was annotated using GeSeq (Tillich et al., 2017), and the three linear contigs were visualized using the same tool. Putative recombination was assessed by mapping Illumina short reads across shared inter-chromosomal repeats with minimap2 (Li, 2021). The assembly graph was visualized with Bandage (Wick et al., 2015) to assess contig conformations. Cis- and trans-splicing gene maps were generated with Circos (Krzywinski et al., 2009) using the annotated exon coordinates of mitochondrial genes. The annotated mitochondrial contigs were submitted to GenBank under the accession numbers PV763367, PV763368 and PV763369.

Phylogenetic analyses were conducted using coding gene sequences from mitochondrial genomes of species within the tribe *Mimoseae*, which comprises approximately 3,300 species (Pedersoli et al., 2023). A search of the NCBI GenBank database yielded complete mitogenome sequences for only seven species. Phylogenetic reconstruction was performed using a concatenated alignment of 35 mitochondrial coding genes from these seven species. Additionally, one species from the *Cassia* clade, *Senna tora* (L.) Roxb., was included as an outgroup. The complete coding genes sequences with default parameters were aligned using MAFFT software (Katoh & Standley, 2013). Using the Corrected Akaike Information Criterion (AICc), the optimal nucleotide substitution model (TVM+G4) for sequence evolution was determined through ModelTest-NG on XSEDE (Darriba et al., 2020). Maximum-likelihood (ML) phylogenetic inference was performed using RAxML-HPC BlackBox v8.1.12 (Stamatakis, 2014) with 1,000 non-parametric bootstrap replicates, executed via the CIPRES Science Gateway v3.3 (Miller et al., 2010). Bootstrap support (BS) values were used to assess the robustness of internal nodes in the resulting phylogenetic trees.

## Results

The mitochondrial genome of *Neltuma alba* (GenBank accessions PV763367, PV763368, and PV763369) consists of three linear contigs, totaling 717,588 base pairs in length (Figure 2A), with an overall GC content of 44.8%. The average read mapping depths of the assembled genome obtained from the NGS sequencing datasets in Chr1-PV763367, Chr2-PV763368 and Chr3-PV763369 were 16,066, 15,433 and 13,417, respectively (Figure S1). Genome annotation identified 59 genes, including 36 protein-coding genes (PCGs), 19 tRNA genes, three rRNA genes, and one pseudogene (*rps7*). Among the PCGs, ten contain introns (*nad1, nad2, nad4, nad5, nad7, cox2, rpl2, rps3, ccmFc* and *rps10*). Of these, *nad4, nad7, cox2, rpl2, rps3, ccmFc* and *rps10* are cis-spliced (Figure S2), whereas *nad1, nad2* and *nad5* are trans-spliced (Figure S3). Intron analysis revealed that *nad1, cox2, ccmFc, rpl2, rps3, nad2* and *rps10* each contain a single intron, while *nad4, nad5*, and *nad7* harbor two or more introns. On the other hand, it is worth noting that the *atp9* gene is duplicated, with two functional copies: one on mtChr2 and another on mtChr3 (Figure 2A). Additionally, several tRNA genes, including *trnE-GUC, trnM-CAU*, and *trnfM-CAU*, are present in multiple copies. Phylogenetic inference using maximum-likelihood (ML) based on 35 mitochondrial PCGs (Figure 2B) revealed four well-supported clades. The outgroup species *Senna tora* (*Cassia* clade) formed a distinct lineage. The remaining species grouped within the tribe *Mimoseae*, which comprised three subclades: (i) *Leucaena* (*L. trichandra, L. leucocephala*, BS = 100); (ii) *Acacia* (*A. ligulata, A. confusa*, BS = 100); and (iii) *Neltuma*, with *N. glandulosa* and *N. alba* forming a subclade (BS = 98), and *N. pallida* as a separate lineage (BS = 100).

**Figure 2.**
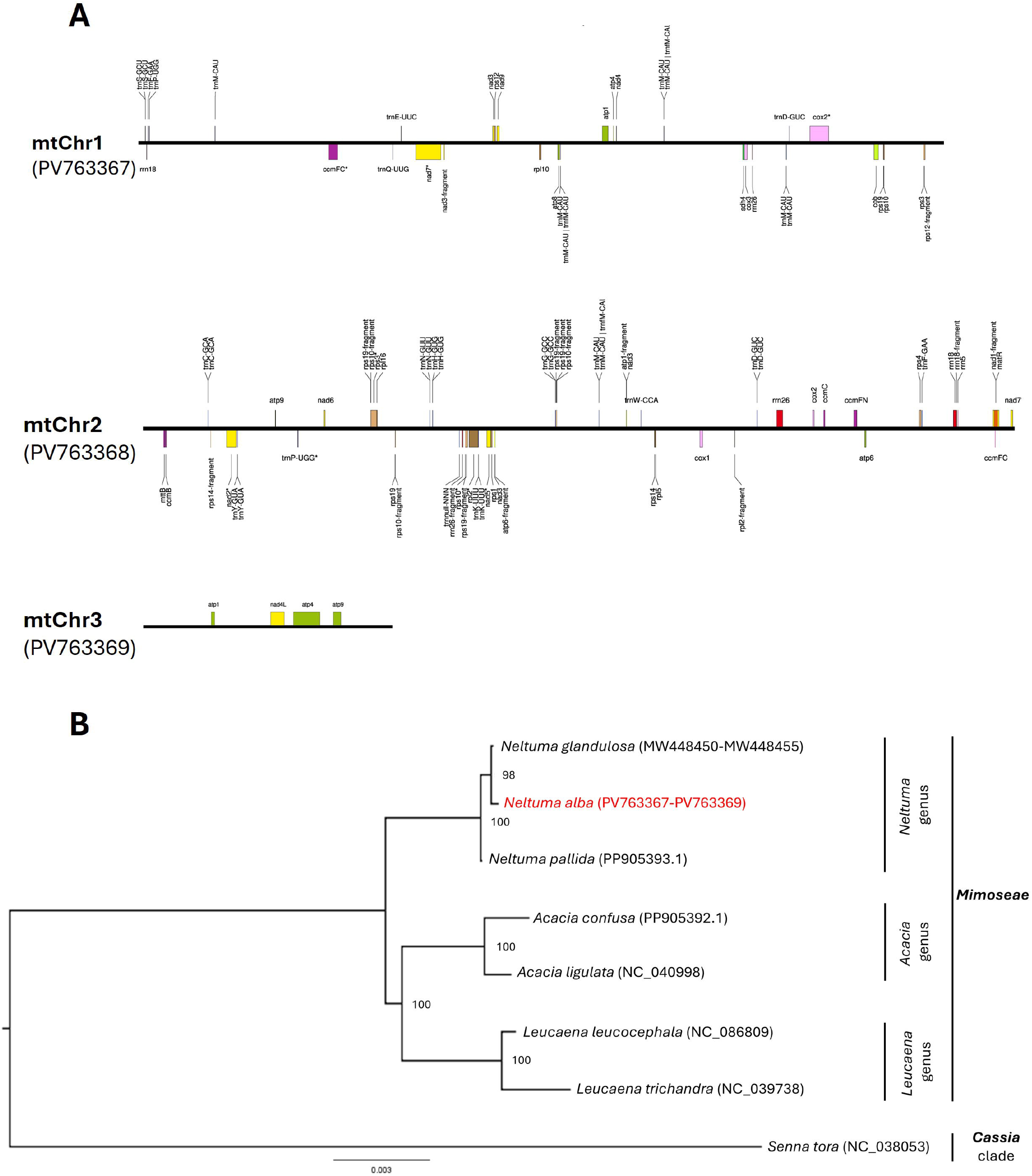
The structural graph of the *Neltuma alba* mitochondrial genome shows three linear chromosomes: mtChr1 with 214,925 base pairs, mtChr2 with 487,804 base pairs, and mtChr3 with 14,859 base pairs. “Gene-fragment” denotes an incomplete gene. (**A**). A maximum-likelihood phylogenetic tree was inferred based on the sequences of 35 mitochondrial coding genes from seven *Mimosoid* clade species (or tribe *Mimoseae*). The following sequences were used: *Neltuma alba* PV763367-PV763369 (synonym *Prosopis alba*) (This study), *Neltuma glandulosa* (Torr.) C.E. Hughes & G.P. Lewis MW448450-MW448455 (synonym *Prosopis glandulosa*) (Choi et al., 2021), *Neltuma pallida* (Humb. & Bonpl. ex Willd.) C.E. Hughes & G.P. Lewis PP905393 (unpublished), *Acacia confusa* Merr. PP905392 (unpublished), *Acacia ligulata* A.Cunn. ex Benth. 1842 NC_040998 (Sanchez-Puerta et al., 2019), *Leucaena leucocephala* (Lam.) de Wit NC_086809 (unpublished), *Leucaena trichandra* (Zucc.) Urb. NC_039738 (Kovar et al., 2018) and *Senna tora* (L.) Roxb. NC_038053 from the *Cassia* clade (Kang et al., 2019), served as the outgroup. The bootstrap support values are placed on the nodes (**B**).

## Discussion and Conclusion

Our assembly reveals 717,588 base pairs distributed across three linear contigs, reflecting the dynamic nature of plant mitogenomes, which frequently exhibit structural plasticity through recombination and multiple configurations (Kozik et al., 2019). Linear mitochondrial chromosomes are expected to be highly sensitive to exonuclease degradation at their termini; therefore, their persistence in vivo implies the action of protective mechanisms. In plant mitochondria, terminal DNA is thought to be shielded within nucleoids and maintained through recombination-dependent repair and replication across shared repeats, which together help stabilize linear configurations and limit deleterious rearrangements (Chevigny et al., 2020; Negroni et al., 2024). Regarding recombination regions, mapping Illumina short reads and analyzing inter-contig repeats revealed 12 candidate regions among the three assembled linear mitochondrial contigs. A closely related species, *Neltuma glandulosa* (syn. *Prosopis glandulosa*), likewise shows a multipartite mitochondrial genome comprising six linear DNA molecules totaling 758,210 base pairs (Choi et al., 2021). Although we used ∼36 Gb of whole-genome Illumina data, the mitochondrial fraction is often low and variable; for context, *Platycodon grandiflorus* achieved 79.1× coverage from 20.9 Gb, whereas *Codonopsis lanceolata* reached 161.4× from just 8.0 Gb (Lee et al., 2018). The annotated mitogenome of *N. alba* includes 36 protein-coding genes (PCGs) and 19 tRNA genes, highlighting conserved functional elements such as the *nad* and *cox* gene families, which are central to stress-responsive energy metabolism (van Aken, 2021). Complementing this, nuclear genome analysis revealed expanded gene families associated with stress tolerance, hormone biosynthesis, photosynthesis, and pathogen resistance—traits indicative of adaptation to drought and salinity (Kong et al., 2023). Although additional mitochondrial genome sequences from tribe *Mimoseae* are available in GenBank, many remain unannotated or unverified, including accessions from the genus *Igna* (OY979714, OY725333, and OY723412). Moreover, some mitogenome sequences used in our phylogenetic analyses - such as PP905393, PP905392, and NC_086809 - are not yet formally published. Phylogenetic reconstruction based on mitogenome genes strongly supports (bootstrap support, BS = 98) the sister-species relationship between *N. alba* and *N. glandulosa*. This placement is consistent with the broader classification of the *Neltuma* clade and aligns with recent taxonomic revisions based on chloroplast genome data (Contreras-Díaz et al., 2024). Furthermore, large-scale phylogenomic analyses using ∼1,000 nuclear genes confirm that the New World *Neltuma* clade, including *N. alba* and *N. glandulosa*, forms a robustly supported monophyletic lineage (Ringelberg et al., 2022). Although mitochondrial data for tribe *Mimoseae* are still scarce, our results emphasize the value of mitogenomic analyses in resolving complex phylogenies and advancing knowledge of desert legume resilience and conservation.

## Authors contributions

Conceptualization: I.R., M.M, M.T, R.C.; Data curation: I.R., M.M, V.R., M.T.; Formal analysis: I.R., M.M, M.T, R.C.; Funding acquisition: R.C.; Investigation: I.R., M.M, R.C.; Methodology: I.R., M.M, V.R., M.T., R.C.; Project administration: R.C.; Resources: R.C.; Software: I.R., M.M, R.C.; Supervision: R.C.; Validation: I.R., M.M, V.R., M.T., R.C.; Visualization: I.R., M.M, R.C.; Writing – original draft: I.R., M.M, V.R., R.C.; Writing – review & editing: I.R., M.M, R.C.

## Ethical approval

This research was conducted in accordance with the required research permits and complies with both national and international standards for the collection of plant material and the protection of flora and fauna. Specifically, collection of *Neltuma alba* (syn. *Prosopis alba*) was authorized by the National Forestry Corporation of Chile (CONAF) under Permit No. N00024/08-11-2019 (JBH/FAP/JVO) and Permit No. N00003-2023/27-01-2023 (NOO/FAP/JVO). Furthermore, this study adheres to the policies of the International Union for Conservation of Nature (IUCN) regarding research involving species at risk of extinction.

## Data availability statement

The data that supports this study is openly available in GenBank of NCBI (https://www.ncbi.nlm.nih.gov) under the accession number PV763367, PV763368 and PV763369. The associated Bio project, Biosample, and SRA numbers are PRJNA1026123, SAMN37734720, and SRR26327158, respectively.

## Supplementary materials

**Figure S1.** Coverage map of the mitogenome assembly.

**Figure S2.** Gene structure map of cis-spliced mitochondrial genes on Chr1 and Chr2 in *Neltuma alba.*

**Figure S3.** Circos map of trans-spliced mitochondrial gene links across Chr1–Chr3 in *Neltuma alba* (*nad1, nad2* and *nad5*).

