## Supplementary figures and images for "Draft mitochondrial genome of the tree species *Neltuma alba* (Caesalpinioideae, tribe *Mimoseae*) from the Atacama Desert, Chile"

### Supplemental Figure 1

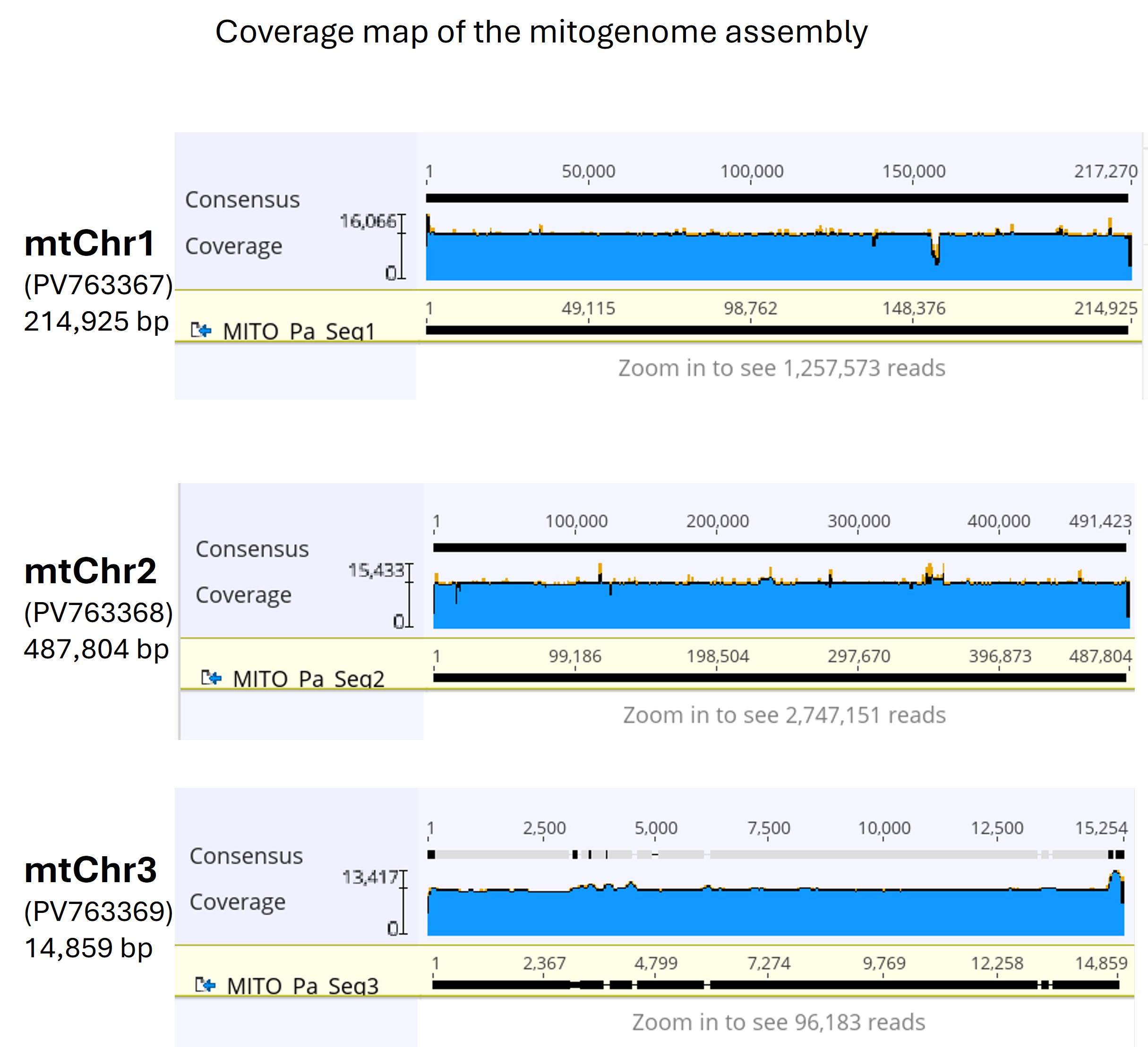

### Supplemental Figure 2

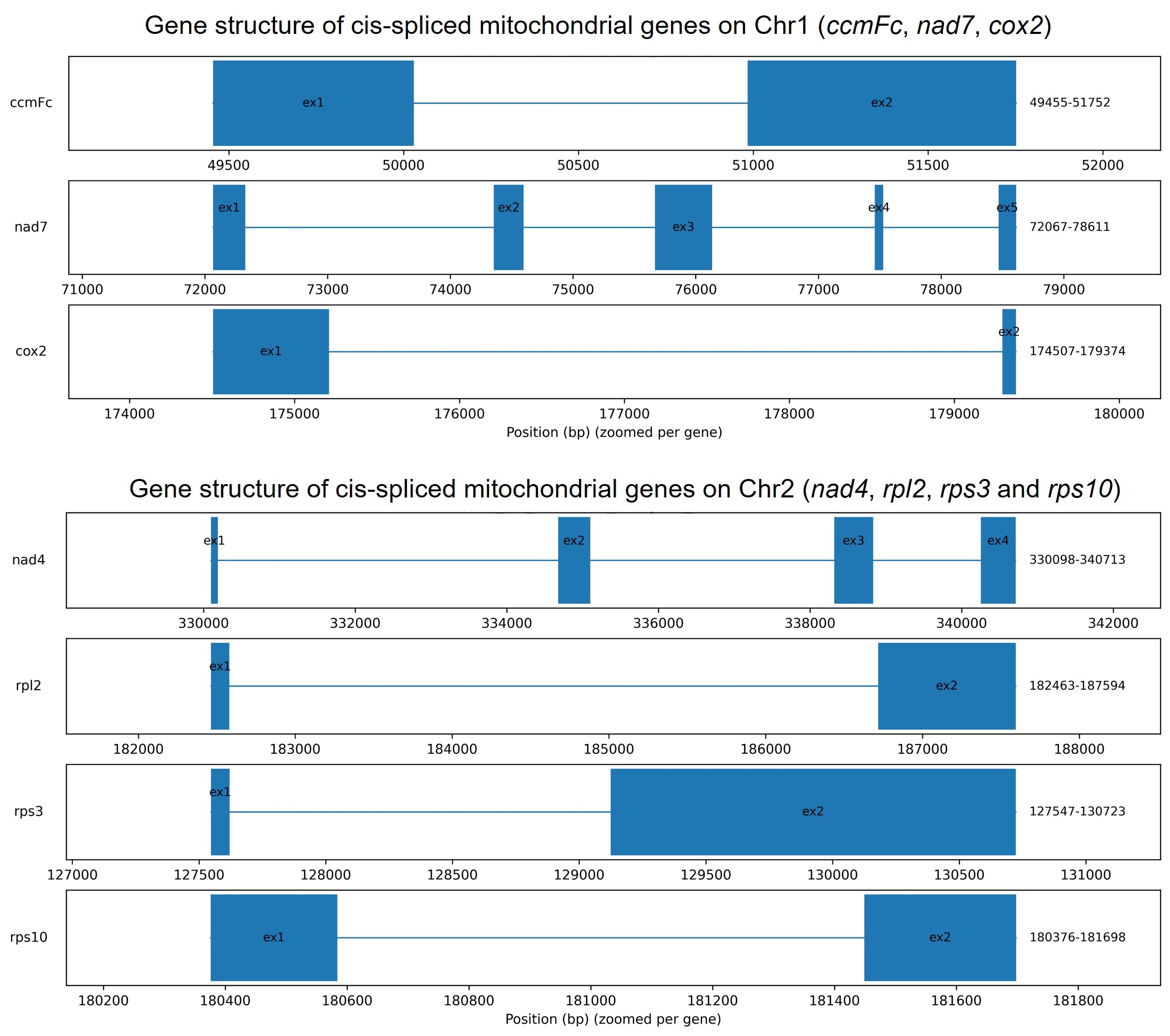

### Supplemental Figure 3

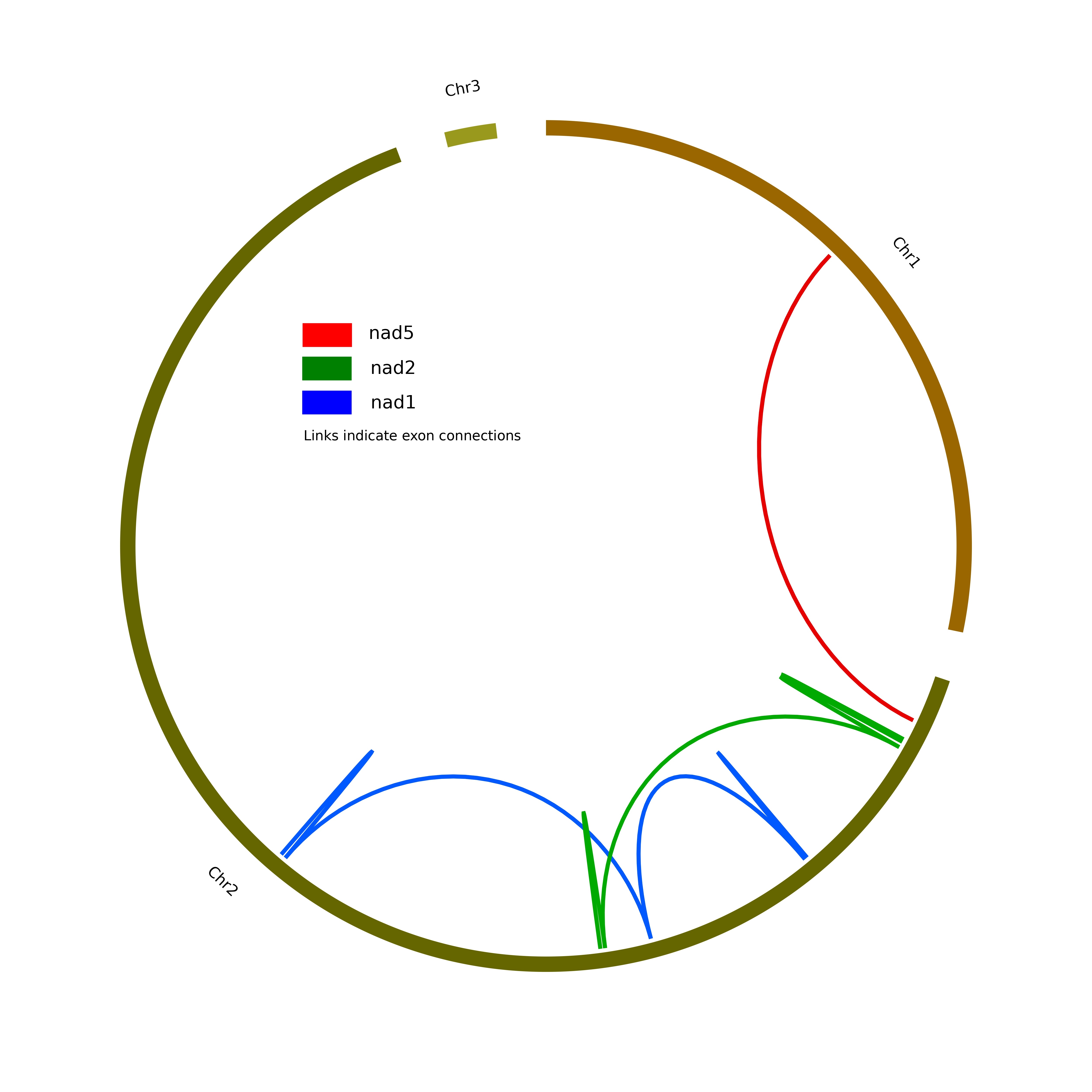
